# Modeling brain dynamics in brain tumor patients using The Virtual Brain

**DOI:** 10.1101/265637

**Authors:** Hannelore Aerts, Michael Schirner, Ben Jeurissen, Dirk Van Roost, Rik Achten, Petra Ritter, Daniele Marinazzo

## Abstract

Presurgical planning for brain tumor resection aims at delineating eloquent tissue in the vicinity of the lesion to spare during surgery. To this end, non-invasive neuroimaging techniques such as functional MRI and diffusion weighted imaging fiber tracking are currently employed. However, taking into account this information is often still insufficient, as the complex non-linear dynamics of the brain impede straightforward prediction of functional outcome after surgical intervention. Large-scale brain network modeling carries the potential to bridge this gap by integrating neuroimaging data with biophysically based models to predict collective brain dynamics.

As a first step in this direction, an appropriate computational model has to be selected, after which suitable model parameter values have to be determined. To this end, we simulated large-scale brain dynamics in 25 human brain tumor patients and 11 human control participants using The Virtual Brain, an open-source neuroinformatics platform. Local and global model parameters of the Reduced Wong-Wang model were individually optimized and compared between brain tumor patients and control subjects. In addition, the relationship between model parameters and structural network topology and cognitive performance was assessed.

Results showed (1) significantly improved prediction accuracy of individual functional connectivity when using individually optimized model parameters; (2) local model parameters can differentiate between regions directly affected by a tumor, regions distant from a tumor, and regions in a healthy brain; and (3) interesting associations between individually optimized model parameters and structural network topology and cognitive performance.

## Introduction

Presurgical planning for brain tumor resection aims at delineating eloquent cortical areas and white matter pathways in the vicinity of the lesion to spare during surgery, in order to preserve essential brain functions. In brain tumor patients, localization of brain function often cannot be inferred from anatomical landmarks alone, since mass effects can distort normal topography or disease processes such as brain shift or plasticity can induce relocation of functions (Desmurget, Bonnetblanc, & Duffau, 2007). Therefore, non-invasive neuroimaging techniques such as functional MRI (fMRI) and diffusion weighted imaging (DWI) fiber tracking are currently employed to pre-operatively determine patient-specific important cortical areas and white matter tracts close to the lesion, to spare during surgery (Sunaert, 2006; Tieleman, Deblaere, Van Roost, Van Damme, & Achten, 2009).

However, taking into account this pre-operative neuroimaging information is often still insufficient, as the complex non-linear dynamics of the brain impede straightforward prediction of functional outcome after surgical intervention. For example, fMRI cannot differentiate cortical areas that are essential for a particular function and which should be surgically preserved, from expendable areas which merely correlate with but are not essential for functionality (Duffau et al., 2003; Tharin & Golby, 2007).

Large-scale brain network modeling carries the potential to bridge this gap by integrating neuroimaging data with biophysically based models to predict collective brain dynamics, from which functionality could be inferred. A future application of this framework could then entail the pre-surgical virtual exploration of the effects of different neurosurgical approaches based on individual patient data, to identify an optimal surgical strategy (Arsiwalla et al., 2015; Proix, Bartolomei, Guye, & Jirsa, 2017).

As a first step in this direction, an appropriate computational model has to be selected, after which suitable model parameter values should be determined that lead to plausible brain dynamics. Different approaches can be used to simulate activity of nodes (i.e., brain areas) in large-scale brain network models varying from detailed spiking neuron models (Deco & Jirsa, 2012), over so-called neural mass or mean-field models that describe the collective activity of cell populations (Deco & Jirsa, 2012), down to abstract models such as the Ising model (Deco, Senden, & Jirsa, 2012; Haimovici, Tagliazucchi, Balenzuela, & Chialvo, 2013; Marinazzo et al., 2014; we refer the interested reader to Deco, Jirsa, Robinson, Breakspear, & Friston (2008) for an in-depth explanation of the general principle behind these approaches and the mechanism allowing a wide class of models to bridge temporal and spatial scales). In a clinical context, biologically interpretable dynamical models are of special interest, as their activity and parameters can allow inference of internal states and processes of the large-scale model, which cannot be measured with non-invasive neuroimaging techniques (Falcon et al., 2015; Schirner, McIntosh, Jirsa, Deco, & Ritter, 2018). Hence, they may provide an entry point for understanding brain disorders at a causal mechanistic level, which might lead to novel, more effective therapeutic interventions (Deco & Kringelbach, 2014; Proix et al., 2017). The first proof-of-concept studies have indicated that global and local biophysical model parameters derived from large-scale brain network modeling can be related to motor recovery after stroke (Falcon et al., 2015) and the generation and progression of epileptic seizures (Bernard & Jirsa, 2017; Jirsa, Stacey, Quilichini, Ivanov, & Bernard, 2014), demonstrating the potential clinical utility of computational modeling.

In this study, we simulated large-scale brain dynamics in brain tumor patients and control subjects using The Virtual Brain (TVB) (Sanz Leon et al., 2013); an open-source neuroinformatics platform that enables the construction, simulation and analysis of brain network models. Local dynamics in each brain region were simulated using the Reduced Wong-Wang model (Deco et al., 2014), one of the more refined and biologically plausible models in the repertoire of TVB. Subsequently, local dynamics of all brain regions were coupled according to each subject’s empirical structural connectome to generate personalized virtual brain models.

The objectives of this study then were (1) to evaluate the importance of constructing personalized virtual brain models using individually optimized model parameters and subject-specific structural connectomes; (2) to determine whether the individually optimized model parameter values differ between brain tumor patients and healthy controls; and (3) to elucidate the relationship between these model parameters on the one hand, and cognitive performance and structural network properties on the other hand.

## Materials and Methods

### Participants

In this study we included patients who were diagnosed with either a glioma, developing from glial cells, or a meningioma, developing in the meninges (Fisher, Schwartzbaum, Wrensch, & Wiemels, 2007). Both types of tumors can be described by their malignancy, based on the World Health Organization grading system. According to this system, grade I tumors are least malignant, whereas grade III (for meningioma) or IV (for glioma) tumors are most malignant. Hereby, malignancy relates to the speed with which the disease evolves, the extent to which the tumor infiltrates healthy brain tissue, and chances of recurrence or progression to higher grades of malignancy.

Patients were recruited from Ghent University Hospital (Belgium) between May 2015 and October 2017. Patients were eligible if they (1) were at least 18 years old, (2) had a supratentorial meningioma (WHO grade I or II) or glioma (WHO grade II or III) brain tumor, (3) were able to complete neuropsychological testing, and (4) were medically approved to undergo MRI investigation. Partners were also asked to participate in the study to constitute a group of control subjects that suffer from comparable emotional distress as the patients.

We collected data from 11 glioma patients (mean age 47.5y, *SD* = 11.3; 4 females), 14 meningioma patients (mean age 60.4y, *SD* = 12.3; 11 females), and 11 healthy partners (mean age 58.6y, *SD* = 10.3; 4 females). Patient characteristics are described in Table 1. Testing took place at Ghent University Hospital (Belgium) on the day before patients’ surgery. All participants received detailed study information and gave written informed consent prior to study enrollment. This research was approved by the Ethics Committee of Ghent University Hospital.

**Table 1.** Patient characteristics. * −1=left-handed, 0=ambidextrous, +1=right-handed; ** (Oligo-)astrocytoma, ependymoma, and oligodendroglioma are subtypes of glioma tumors.

| Subject ID | Sex | Age | Handed-ness* | Tumor lateralization | Tumor location | Tumor size (cm<sup>3</sup>) | Tumor histology** |
| --- | --- | --- | --- | --- | --- | --- | --- |
| PAT01 | F | 67 | 1.00 | Left | Frontal | 1.69 | Meningioma I |
| PAT02 | F | 49 | 1.00 | Right | Frontal | 0.75 | Meningioma I |
| PAT03 | F | 60 | 1.00 | Right | Parietal | 78.44 | Meningioma I |
| PAT05 | F | 40 | 1.00 | Left | Frontal | 12.95 | Oligo-astrocytoma II |
| PAT06 | F | 67 | 1.00 | Bilateral | Frontal | 12.81 | Meningioma I |
| PAT07 | M | 55 | -0.16 | Left | Temporal | 33.94 | Ependymoma II |
| PAT08 | M | 51 | 1.00 | Left | Frontal | 16.81 | Meningioma I |
| PAT10 | F | 77 | 1.00 | Left | Frontal | 0.59 | Meningioma I |
| PAT11 | F | 74 | 1.00 | Left | Frontal | 20.73 | Meningioma II |
| PAT13 | F | 40 | -0.68 | Bilateral | Skull base | 2.09 | Meningioma I |
| PAT14 | F | 74 | 1.00 | Left | Parietal | 3.45 | Meningioma I |
| PAT15 | F | 57 | 1.00 | Right | Frontal | 2.13 | Meningioma I |
| PAT16 | M | 39 | 1.00 | Right | Fronto-temporal | 50.24 | Anaplastic astrocytoma II-III |
| PAT17 | F | 49 | 1.00 | Right | Frontal | 0.58 | Meningioma I |
| PAT19 | F | 44 | 1.00 | Right | Frontal skull base | 2.64 | Meningioma I |
| PAT20 | F | 70 | 0.60 | Right | Parietal | 15.24 | Anaplastic astrocytoma III |
| PAT22 | M | 53 | 1.00 | Right | Parietal | 15.16 | Oligodendroglioma II |
| PAT23 | M | 74 | 1.00 | Bilateral | Frontal | 89.56 | Meningioma I |
| PAT24 | M | 62 | 1.00 | Left | Frontal | 5.91 | Meningioma I |
| PAT25 | F | 46 | 1.00 | Right | Temporal | 18.56 | Glioma II |
| PAT26 | M | 61 | 1.00 | Right | Temporal | 59.21 | Anaplastic astrocytoma III |
| PAT27 | M | 38 | 1.00 | Left | Frontal | 13.24 | Astrocytoma II |
| PAT28 | M | 44 | 1.00 | Left | Frontal | 11.49 | Oligodendroglioma II |
| PAT29 | M | 45 | 0.90 | Left | Frontal | 36.20 | Oligodendroglioma III |
| PAT31 | F | 31 | 0.90 | Left | Frontal | 4.24 | Oligodendroglioma II |

### MRI data acquisition

From all participants, three types of MRI scans were obtained using a Siemens 3T Magnetom Trio MRI scanner with a 32-channel head coil. First, T1-MPRAGE anatomical images were acquired (160 slices, *TR* = 1750 ms, *TE* = 4.18 ms, field of view = 256 mm, flip angle = 9°, voxel size 1 × 1 × 1 mm, *TA* = 4:05 min). Next, resting-state functional echo-planar imaging (EPI) data were obtained in an interleaved order (42 slices, *TR* = 2100ms, *TE* = 27 ms, field of view = 192 mm, flip angle = 90°, voxel size 3 × 3 × 3 mm, *TA* = 6:24 min). After the first 4 control subjects, 5 meningioma patients and 2 glioma patients were scanned, the fMRI protocol was accidentally changed to a *TR* of 2400 ms resulting in a *TA* of 7:19 minutes. This has been taken care of in subsequent analyses by inclusion of an additional covariate. During the fMRI scan, participants were instructed to keep their eyes closed and not fall asleep. Finally, a multi-shell high-angular resolution diffusion-weighted MRI (DWI) scan was acquired (60 slices, *TR* = 8700 ms, *TE* = 110 ms, field of view = 240 mm, 101 diffusion directions, b-values of 0, 700, 1200, 2800 s/mm^2^, voxel size 2.5 × 2.5 × 2.5 mm, *TA* = 15:14 min). In addition, two DWI b = 0 s/mm2 images were collected with reversed phase-encoding blips for the purpose of correcting susceptibility induced distortions (Andersson, Skare, & Ashburner, 2003).

### MRI data preprocessing

MRI data were preprocessed and subject-specific structural and functional connectivity matrices were extracted using a modified version of the TVB preprocessing pipeline (Schirner, Rothmeier, Jirsa, McIntosh, & Ritter, 2015). All steps are outlined below.

#### Preprocessing of T1-weighted anatomical MRI data

In the first step, high-resolution anatomical images were processed using FreeSurfer (http://surfer.nmr.mgh.harvard.edu) to obtain a subject-specific parcellation of each subject’s brain into 68 cortical regions (34 per hemisphere). T1-weighted data of all control subjects were subjected to the default recon-all processing pipeline, which includes the following steps: intensity normalization, skull stripping, removal of non-brain tissue, brain mask generation, cortical reconstruction, segmentation of subcortical white matter and deep gray matter volumetric structures, cortical tessellation of the gray matter/white matter and gray matter/pial boundary, and construction of a probabilistic atlas based cortical parcellation into 68 regions according to gyral and sulcal structure (Desikan et al., 2006; Fischl et al., 2004).

As meningioma tumors generally exert pressure on the brain without infiltrating, our aim was to segment out the meningioma tumor before cortical reconstruction. However, visual inspection of the results showed this was done automatically by the recon-all processing pipeline of FreeSurfer in all but two meningioma patients. In the remaining two meningioma patients, who had very large lesions, manual edits were made.

Glioma tumors, in contrast, generally do infiltrate the brain. To obtain a whole-brain parcellation scheme for these patients, two additional steps were carried out. First, glioma tumors were segmented using the Unified Segmentation with Lesion toolbox (https://github.com/CyclotronResearchCentre/USwithLesion, unpublished observations). Second, the Normalisation tool of the BCBtoolkit (Foulon et al., 2018) was used to produce an enantiomorphic filling of the affected area by symmetrically filling up the lesion mask with healthy tissue of the contralateral hemisphere (Nachev, Coulthard, Jäger, Kennard, & Husain, 2008). These “normalized” anatomical MRI data were then processed using the standard recon-all FreeSurfer processing pipeline. Resulting parcellations were visually inspected and manually corrected in two glioma patients.

#### Functional MRI preprocessing

Functional MRI (fMRI) data processing was carried out using FEAT (FMRI Expert Analysis Tool, version 6.00), part of FSL (FMRIB’s Software Library, http://www.fmrib.ox.ac.uk/fsl). Specifically, the following operations were applied: motion correction using MCFLIRT (Jenkinson, Bannister, Brady, & Smith, 2002), slice-timing correction, non-brain removal using BET (S. M. Smith, 2002), grand-mean intensity normalization of the entire 4D dataset by a single multiplicative factor, and high-pass temporal filtering (100-second high-pass filter). Next, the FreeSurfer cortical parcellation obtained in the previous step was mapped to the subject’s functional space. To this end, fMRI images were linearly registered to the subject’s high-resolution T1-weighted images using the epi_reg function of FSL FLIRT (Jenkinson et al., 2002; Jenkinson & Smith, 2001), after which the inverse of this transformation matrix was applied to transform the FreeSurfer parcellation scheme to the subject’s functional space. Average BOLD signal time series for each region were then generated by computing the spatial mean for all voxel time-series of each region. Lastly, functional connectivity (FC) matrices were constructed by calculating the Fisher z-transformed Pearson correlation coefficient between all pairs of region-wise aggregated BOLD time series.

#### Diffusion-weighted MRI preprocessing

Since all analyses of this study depend on the quality of the structural connectivity matrices, a state-of-the art pipeline was constructed for the preprocessing of DWI data and consecutive network construction, using a combination of FSL (FMRIB’s Software Library, http://www.fmrib.ox.ac.uk/fsl; version 5.0.9) and MRtrix3 (http://www.mrtrix.org; version 0.3.RC2). First, raw diffusion weighted MRI images were corrected for several artifacts. In particular, DWI images were denoised (MRtrix dwidenoise; Veraart et al., 2016), corrected for Gibbs ringing artifacts (MRtrix mrdegibbs; Kellner, Dhital, & Reisert, 2016), for motion and eddy currents (FSL eddy; Andersson & Sotiropoulos, 2016), for susceptibility induced distortions (FSL topup; Andersson, Skare, & Ashburner, 2003), and for bias field inhomogeneities (FSL FAST; Zhang, Brady, & Smith, 2001). Next, subjects’ high-resolution anatomical images were linearly registered to diffusion space with the epi_reg function of FSL FLIRT (Jenkinson et al., 2002; Jenkinson & Smith, 2001), and subsequently segmented into gray matter, white matter and cerebrospinal fluid (FSL FAST; Zhang, Brady, & Smith, 2001).

Then, DWI images were intensity normalized across subjects and group average response functions were calculated. Specifically, response functions for each subject were estimated per b-value shell (b = 0, 700, 1200 & 2800 s/mm^2^) and per tissue type (white matter, gray matter, cerebrospinal fluid) using the MRtrix3 script dwi2response dhollander (Dhollander, Raffelt, & Connelly, 2016). A scaling factor per subject was calculated by which the individual response functions could be multiplied in order to obtain the average response function across all subjects. DWI images were then initially normalized by dividing subjects’ DWI images by their corresponding scaling factor. After that, response functions per b-value shell and tissue type were recalculated for every subject and averaged across all subjects. This set of group average response functions were subsequently utilized in multi-shell multi-tissue constrained spherical deconvolution to estimate the fiber orientation distributions (MRtrix3 msdwi2fod; Jeurissen, Tournier, Dhollander, Connelly, & Sijbers, 2014). In addition, tissue components from multi-tissue CSD were once more intensity normalized using MRtrix3 mtnormalise.

Next, anatomically constrained probabilistic whole-brain fiber tracking (ACT) was performed using dynamic seeding generating 30 million streamlines per subject (MRtrix3 tckgen; R. E. Smith, Tournier, Calamante, & Connelly, 2015, 2012). Afterwards, spherical-deconvolution informed filtering of tractograms (SIFT) was applied to selectively filter out streamlines from the tractogram in order to improve the fit between the streamline reconstruction and the underlying diffusion images, retaining 7.5 million streamlines per subject (MRtrix3 sift; R. E. Smith, Tournier, Calamante, & Connelly, 2013). A structural connectivity (SC) matrix was then constructed by transforming the individual’s FreeSurfer parcellation scheme to diffusion space, and calculating the number of estimated tracts between any two brain regions (MRtrix3 tck2connectome). In addition, a distance matrix was constructed by calculating the average length of all streamlines connecting any two nodes (MRtrix3 tck2connectome). By using a proper high-order model and taking into account the full fiber orientation distribution function (through CSD), by taking into account the presence of non-white matter tissue (through multi-tissue CSD), by applying realistic individual anatomical priors (through ACT), and by ensuring fidelity of the tractograms to the data (through SIFT), it has been shown that the biological accuracy of tractograms can be vastly increased compared to those obtained with unfiltered unconstrained diffusion tensor tracking (Jeurissen, Descoteaux, Mori, & Leemans, 2017).

Lastly, we thresholded and normalized the resulting SC matrices. Thresholding was carried out to minimize false-positive streamlines. Using an absolute threshold, setting to zero all connection weights smaller than 5, yielded a decaying degree distribution while ensuring all subjects’ network remained fully connected. This approach is similar to the one adopted by Collin, Kahn, De Reus, Cahn, & Van den Heuvel (2014). Normalization was performed by dividing all SC weights by a constant scalar across subjects (75.000 in our case: 7.5 million streamlines generated per subject / 100) to ensure all SC weights varied between 0 and 1, which was required for computational modeling in TVB.

### Computational modeling

TVB (Sanz Leon et al., 2013) was utilized to simulate large-scale brain dynamics (see Figure 1 for visual overview of the workflow). In particular, local dynamics in each of the 68 cortical brain regions were simulated using the Reduced Wong-Wang model (Deco et al., 2014), implemented as highly optimized C code that allows for efficient parameter exploration (Schirner et al., 2018). In particular, each cortical region of the Desikan-Killiany brain atlas (Desikan et al., 2006) was modeled as a local network composed of interconnected excitatory and inhibitory neural mass models coupled by excitatory (NMDA) and inhibitory (GABA) connections (we refer the interested reader to Deco et al. (2014) for a more detailed description of this model). Excitatory neural mass models of all 68 brain regions were subsequently coupled according to the individual subject’s empirical structural connectome and weighted by a global scaling factor, to simulate large-scale brain dynamics.

**Figure 1.**
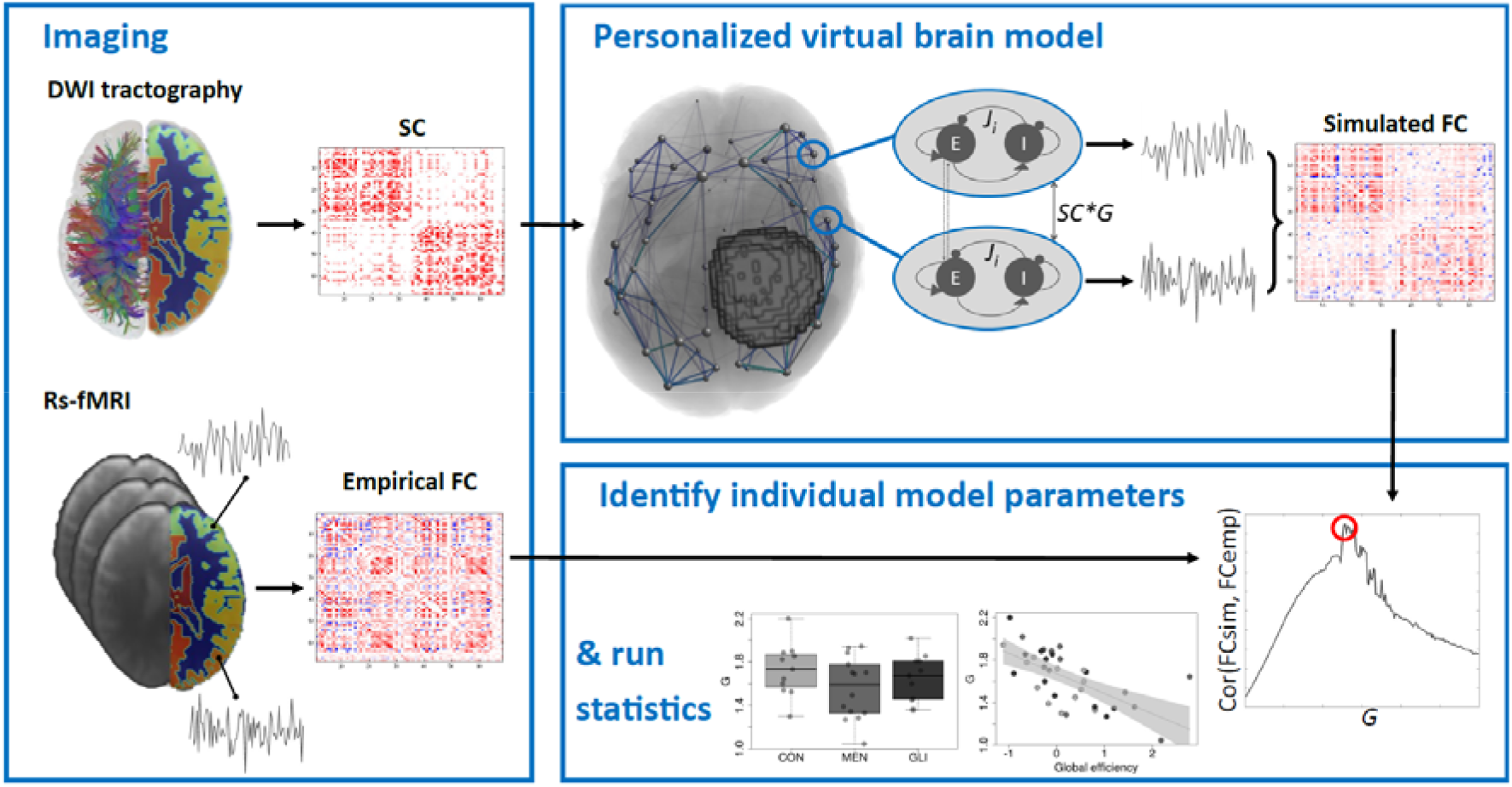
Graphical summary of computational modeling workflow using TVB. **Imaging**: As input to the model, individual neuroimaging data should be acquired. Diffusion MRI data processed to a structural connectivity (SC) matrix as input for TVB, and resting-state functional MRI data processed to an empirical functional connectivity (FCemp) matrix to evaluate model fit (red=positive connection weights, blue=negative connection weights). **Personalized virtual brain model**: Local dynamics in each of the 68 Freesurfer cortical brain regions were simulated using the Reduced Wong-Wang model (Deco et al., 2014), implemented as highly optimized C code that allows for efficient parameter exploration (Schirner et al., 2018). Excitatory neural mass models of all 68 brain regions were subsequently coupled according to the individual subject’s empirical structural connectome and weighted by a global scaling factor (*G*), to simulate large-scale brain dynamics. Resting-state blood-oxygen-level-dependent (BOLD) time series were generated with the same duration and sampling rate as the subject’s empirical resting-state fMRI acquisition, using the Balloon-Windkessel hemodynamic model (Friston et al., 2000). From these simulated time series, a simulated functional connectivity (FCsim) matrix was computed. **Identify optimal model parameters**: To optimize the fit between empirical and simulated FC, subject-specific parameter space explorations were conducted in which the global scaling parameter (*G*) was optimized. For each value of the global scaling parameter, a simulated FC matrix was constructed. The value of the global scaling parameter that maximized the link-wise Pearson correlation between each individual’s simulated and empirical functional connectivity matrix was then selected for further analyses. **& run statistics**: Individually optimized model parameters were subsequently tested for group differences and correlated with structural graph theory metrics and cognitive performance.

In order to optimize the fit between empirical and simulated functional connectivity, subject-specific parameter space explorations were conducted in which the global scaling parameter (*G*) was optimized. This parameter rescales each subject’s structural connectivity, which is given by relative values, to yield absolute interaction strengths. In particular, values of the global scaling parameter were varied from 0.01 to 3 in steps of 0.015. For each value of the global scaling parameter, individual resting-state blood-oxygen-level-dependent (BOLD) time series were generated with the same duration and sampling rate as the subject’s empirical resting-state fMRI acquisition, using the Balloon-Windkessel hemodynamic model (Friston, Mechelli, Turner, & Price, 2000). From these simulated time series, a functional connectivity matrix was computed by calculating the Fisher z-transformed Pearson correlation coefficient between all pairs of simulated BOLD time series. The value of the global scaling parameter that maximized the link-wise Pearson correlation between each individual’s simulated and empirical functional connectivity matrix was then selected for further analyses.

In addition, the feedback inhibition control parameters (*J_i_*) – controlling the strengths of connections from inhibitory to excitatory mass models within each large-scale region *i* – were tuned in each iteration of the parameter space exploration, to clamp the average firing rate at ~3 Hz for each excitatory mass model. 3 Hz was chosen as attractor value, as it is the “intrinsic frequency” of an isolated neural mass model according to derivations from a spiking network (Deco et al., 2014). When multiple mass models are coupled, as in a virtual brain model, the firing rates increase due to the input from the global network. To get the firing rates back to a physiologically plausible average rate of 3 Hz, the feedback inhibition control parameter *J_i_* is increased until the population has this average firing rate. Thus, 3 Hz is the attractor value for each population, around which there is an ongoing fluctuation during the entire simulation time. Previous work has shown that tuning of these local model parameters significantly improves prediction of empirical functional connectivity (Deco et al., 2014). Note that only the global coupling scaling factor was used to maximize the fit with empirical functional connectivity, while the sole fitting target of the local inhibitory connection strengths were the average firing rates of excitatory mass models.

Given the skewed distribution of local inhibitory connection strengths, median *J_i_* values were computed per subject for further analyses. First, the median across the entire brain was computed (*J_brain_*). Second, median *J_i_* was calculated in patients across a subset of tumor regions (*J_tumor_*) to investigate possible local alterations in biophysical model parameters in the direct vicinity of the lesion. In glioma patients, tumor regions were defined as those cortical areas of the individual FreeSurfer parcellation that showed at least partial (i.e. minimum 1 voxel) overlap with the tumor mask. In meningioma patients, tumor regions consisted of regions that were (at least partially) displaced because of the tumor’s mass effect. To estimate which regions were displaced by the meningioma, patients’ anatomical images were transformed to MNI space (using FSL FLIRT with 12 DOF), and this transformation was applied to their tumor mask. Then, the overlap between subjects’ tumor mask in MNI space and the fsaverage Desikan-Killiany atlas (Desikan et al., 2006) in MNI space was calculated. Parcels that showed at least 1 voxel overlap with the tumor mask were denoted tumor nodes. Table 1-1 gives a graphical overview of the extent to which tumor nodes were affected in meningioma and glioma patients. Lastly, we also computed median local inhibitory connection strengths across healthy regions (*J_non-brain_*) to investigate possible distant lesion effects. In control subjects, this was the same as the whole-brain median *J_i_*, whereas in tumor patients, the median across all non-tumor regions was calculated.

**Table 1-1.**
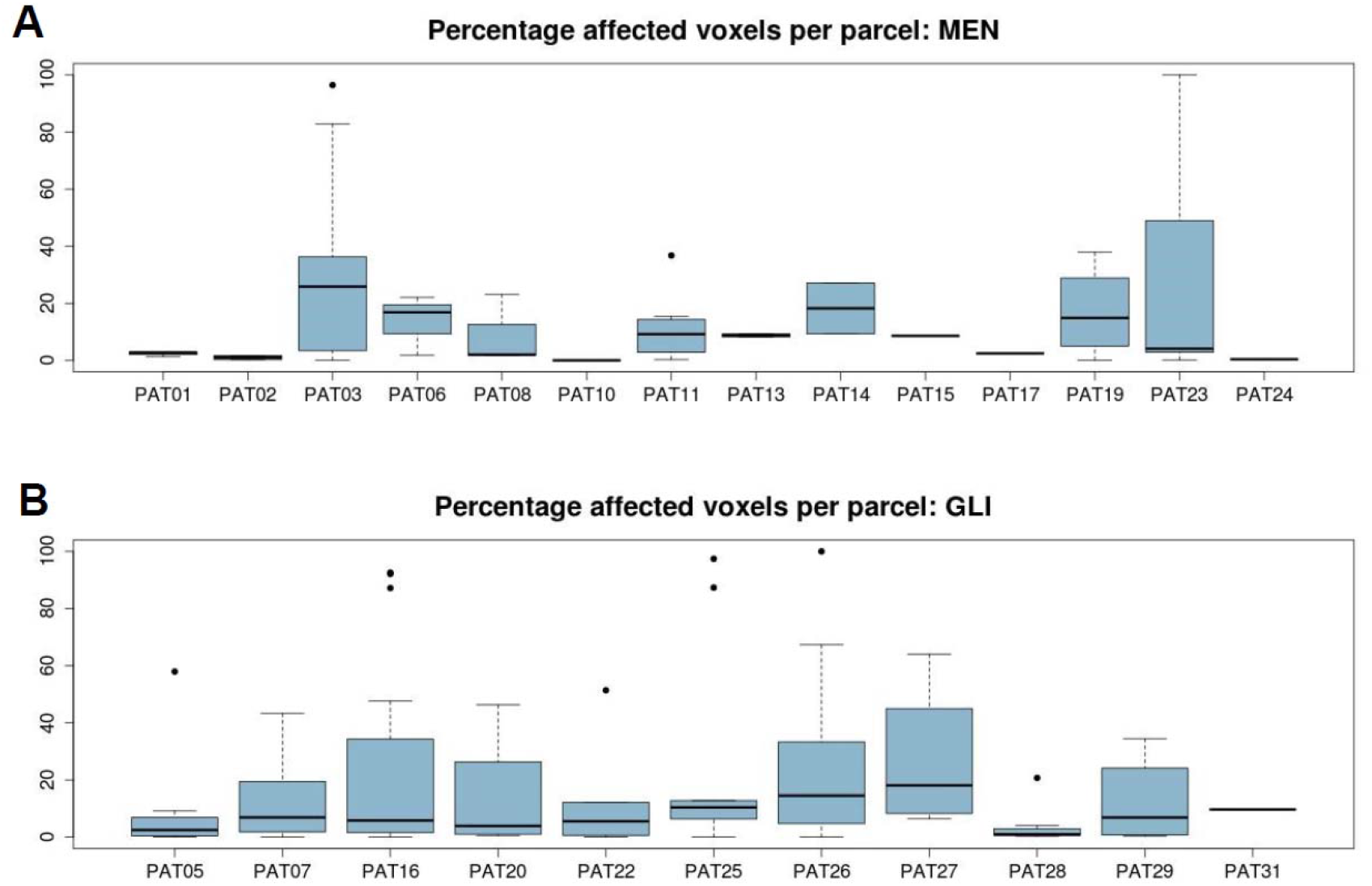
Percentage of voxels per node that was affected by a tumor (only for tumor nodes), per subject, for (A) meningioma patients and (B) glioma patients.

### Graph theory analysis

Structural network topology was assessed with various graph theory metrics of integration (global efficiency, communicability), segregation (clustering coefficient, local efficiency, modularity), and centrality (degree, strength, betweenness centrality, participation coefficient), as well as graph density, using the Brain Connectivity Toolbox (Rubinov & Sporns, 2010). After inspecting the relationships among these graph theory metrics by linear correlation and principal component analysis, three distinct graph theory metrics were retained (|*r*|<0.80 and graph metrics separable by the first 2 principal components) for further analyses: global efficiency, modularity, and participation coefficient.

Global efficiency relates to the capacity of the network to rapidly integrate specialized information from distributed brain regions, and is defined as the average inverse shortest path length (Latora & Marchiori, 2001). Modularity, on the other hand, is a measure of segregation, estimating the size and composition of densely interconnected groups of nodes. The modular structure can be revealed by subdividing the network into modules by maximizing the number of within-group links and minimizing the number of between-group links (Girvan & Newman, 2002; Guimerà & Amaral, 2005). Participation coefficient then measures the centrality of a node by the ratio of its intramodular to intermodular connections (Guimerà, Sales-Pardo, & Amaral, 2007). Of note, the participation coefficient and modularity index Q were both calculated using the same modular decomposition, which was identified through modularity maximization across 100 iterations. As an additional check, we calculated the stability of this modular decomposition across iterations per subject. In particular, we calculated (per subject) how often any 2 nodes were grouped within the same module. Then, we computed the average stability across all nodes. Results showed relatively high stability (average across subjects = 90.25%, *SD* = 7.47%). For more details and an in-depth discussion of graph metrics, we refer the interested reader to Rubinov & Sporns (2010).

### Neuropsychological testing

Cognitive performance of patients and control subjects was assessed using the Cambridge Neuropsychological Test Automated Battery (CANTAB®; Cambridge Cognition (2017); All rights reserved; http://www.cambridgecognition.com). In particular, four cognitive domains were examined that have been identified by previous studies to be affected by brain tumors: sustained attention, working memory, information processing speed, and executive functioning (Derks, Reijneveld, & Douw, 2014). One glioma patient was not tested because of lack of time, and one meningioma patient did not complete the sustained attention task.

Prior to the actual test administration, the Motor Screening Task (MOT) was used to screen for sensorimotor or comprehension difficulties that could limit valid data collection. Subsequently, the main tasks were administered in random order to avoid sequence bias. Specifically, the Rapid Visual Information Processing (RVP) task was used to assess sustained attention, the Spatial Span (SSP) task measured working memory capacity, the Reaction Time task (RTI) evaluated participants’ mental response speed, and the Stockings of Cambridge (SOC) task assessed planning accuracy.

### Accounting for covariates of no immediate interest

As cognitive performance can be affected by several factors, results were corrected for participants’ motivation, level of emotional distress, lesion volume, age and sex. Likewise, previous studies have shown that graph theory metrics can be affected by various factors such as age and gender (Biswal et al., 2010), handedness (Bettus et al., 2010), and mood (Harrison et al., 2008). Therefore, graph theory results were corrected for these confounding variables, as well as for lesion volume and several MRI parameters (motion during rs-fMRI acquisition, TR of rs-fMRI protocol, intensity normalization factor used in DWI preprocessing). In particular, we used the inverse of the mean latency with which participants responded to the MOT task as a proxy of their motivation. To measure emotional distress/mood on the day testing took place, the Dutch version of the State-Trait Anxiety Inventory (Spielberger, Gorsuch, Lushene, Vagg, & Jacobs, 1983; van der Ploeg, 1982) was utilized. Further, lesion volume was calculated as the number of 1 mm^3^ isotropic voxels in the tumor mask drawn on the anatomical T1 image, and handedness was measured using the Edinburgh Handedness Inventory (Oldfield, 1971).

Linear regression models were then constructed for every outcome variable (sustained attention, working memory capacity, mental reaction time, and planning accuracy for cognitive performance; global efficiency, modularity and participation coefficient for graph theory metrics) as a function of these confounders. Of note, only main effects of the confounders were considered, since inclusion of interaction effects between the covariates would considerably reduce the degrees of freedom. Residuals of these models were further transformed to z-scores for subsequent analyses using the mean and standard deviation of the respective metric in the group of control subjects, for the ease of interpretation.

Additionally, we investigated the effect of TR on the construction of FC matrices and the parameter space exploration. First, we simulated BOLD timeseries for every subject, using the subject’s optimized global coupling value and (A) a TR corresponding to the subject’s empirical rs-fMRI TR, and (B) the other TR (i.e., for a subject who was scanned with rs-fMRI TR = 2.1, we then used TR = 2.4 and vice versa). Results showed that the similarity between the upper triangular part of both simulated FC matrices per subject was rather high (average Pearson correlation across subject = 0.97, *SD* = 0.025). Further, we investigated in two subjects (one control subject and one glioma patient) whether the parameter space exploration was comparable when using a different TR (i.e. using TR = 2.1 while the original rs-fMRI TR for these subjects was 2.4). Results again showed a very limited impact of TR on the parameter space exploration.

### Statistical analyses

First, we compared the optimized model parameters between glioma patients, meningioma patients and control participants using one-way analysis of variance tests and Kruskal-Wallis rank sum tests, depending on whether or not the normality assumption was violated. Afterwards, optimal model parameters were related to structural network topology and cognitive performance using linear regression. All analyses were performed on a Cooler Master Ubuntu 14.04 desktop pc.

## Results

### Model fit of personalized virtual brain models

In this study, we simulated large-scale brain dynamics in brain tumor patients and control subjects using the Reduced Wong-Wang model (Deco et al., 2014) as implemented in a highly optimized C version of TVB’s simulation core (Schirner et al., 2018). After tuning the model parameters, the link-wise Pearson correlation between the upper triangular part of the simulated and empirical functional connectivity matrices was on average 0.33 across subjects (*SD* = 0.09). These magnitudes of similarity are similar to those reported in other studies that have simulated subject-specific brain dynamics at the large-scale level (see for example Deco et al., 2013; Schirner et al., 2018). The obtained correlations between simulated and empirical functional connectivity in turn correlated with the correlation coefficients between the subject’s structural and empirical functional connectivity (Pearson’s *r*(SC-FC_emp_, FC_emp_-FC_sim_) = 0.68, 95% CI 0.45-0.82, *p* < 0.0001). No significant differences in prediction accuracy of empirical functional connectivity were found between healthy controls, meningioma patients and glioma patients (*F*(2,33) = 0.32, *p* = 0.73) or according to lesion size (*F*(1,34) = 1.20, *p* = 0.28).

Additional analyses demonstrated that individual optimization of the modeling parameters significantly contributed to the prediction of subjects’ FC (ANOVA F(3,140) = 6.34, *p* = 0.0005) (Figure 2). In particular, post-hoc Tukey tests showed that highest similarity between individuals’ simulated and empirical FC was obtained when model parameters were individually tuned on the individual or control average SC matrix *(diff* = 0.007, *p* = 0.984 for “Indiv SC; Indiv params, optimized on indiv SC” [Fig 2D] vs. “CON avg SC; Indiv params, optimized on CON avg SC” [Fig 2C]). When simulating FC using individual structural connectomes with average model parameters from the control group [Fig 2A], or using a control average SC matrix with individual model parameters obtained from parameter space explorations on the individual SC matrices [Fig 2B], prediction accuracies were significantly lower (*diff* = 0.06, *p* = 0.010 for “Indiv SC; Indiv params, optimized on indiv SC” [Fig 2D] vs. “Indiv SC, CON avg params” [Fig 2A]; *diff* = 0.07, *p* = 0.006 for “Indiv SC; Indiv params, optimized on indiv SC” [Fig 2D] vs. “CON avg SC; Indiv params, optimized on indiv SC” [Fig 2B]).

**Figure 2.**
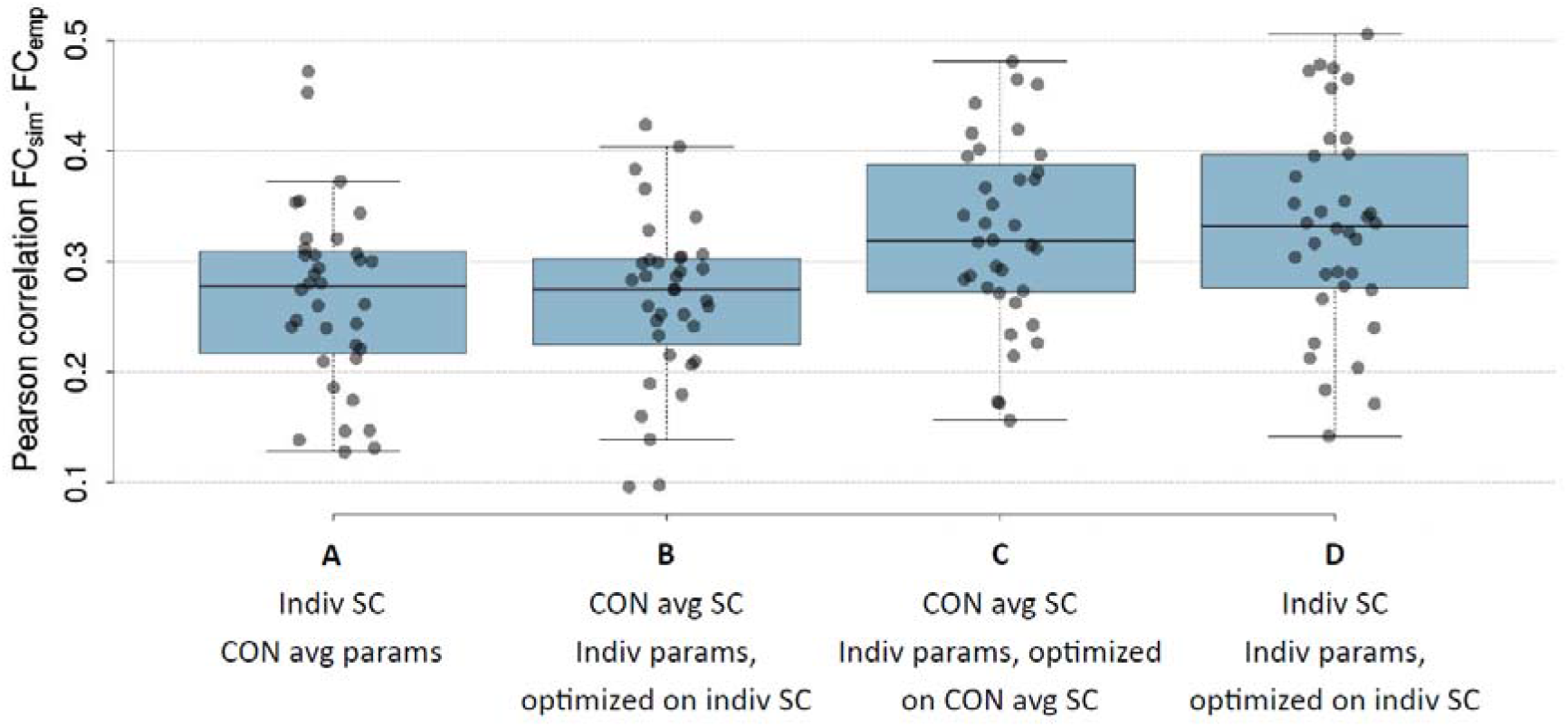
Pearson correlation coefficient between simulated and empirical FC for all 36 subjects. Simulated FC was obtained using (1) average model parameters of control group with individual SC matrices; (2) control average SC matrix with individually optimized model parameters obtained from parameter space explorations on the individual SC matrices; (3) control average SC matrix with individually optimized model parameters obtained from parameter space explorations on the control average SC matrix; and (4) individually optimized model parameters and individual SC matrices.

### Local model parameters altered in brain tumor patients

We then sought to determine whether the individually optimized biophysical model parameter values differ between glioma patients, meningioma patients and control participants. Figure 3A shows the distribution of optimal values of the local inhibitory connection strengths across the entire brain of healthy control subjects, and across tumor regions in meningioma and glioma patients. Results revealed that median local inhibitory connection strengths were marginally more variable in tumor regions compared to healthy brains (Levene’s test of equality of variances *F*(2,33) = 3.44, *p* = 0.044). Furthermore, a trend towards increased group means in tumor regions was observed, although this difference did not reach statistical significance (ANOVA *F*(2,33) = 2.72, *p* = 0.081). In healthy brain regions, median local inhibitory connection weights did not differ between patients and controls (Figure 3B) (ANOVA *F*(2,33) = 1.27, *p* = 0.293). However, local inhibitory connection strengths strongly depend on the number of connections a brain region has, which in turn tightly correlates with region size. That is, larger cortical areas encompass on average more white matter fiber bundles (i.e., higher in-/out-strength), and hence need more local inhibition to balance global excitation. Additional analyses showed that tumor regions were significantly larger compared to non-tumor regions (Kruskal-Wallis *X^2^*(1) = 15.11, *p* = 0.0001), but that the number of connections per cortical area were not significantly different in tumor regions compared to non-tumor regions (*X^2^*(1) = 2.29, *p* = 0.130). Given the high dependency between region in-strength and size (*r* = 0.80), we therefore only regressed out the effect of region size. Results now revealed opposite effects (Figure 3C), with median local inhibitory connection strengths that are much lower (Kruskal-Wallis *X^2^*(2) = 14.4, *p* = 0.0007) and more variable (Levene’s test of equality of variances *F*(2,33) = 5.27, *p* = 0.010) in tumor regions compared to healthy regions/brains. Moreover, local inhibitory connection strengths in non-tumor regions were now also increased (Kruskal-Wallis *X^2^*) = 6.51, *p* = 0.039) compared to healthy controls (Figure 3D). With regard to the global coupling parameter (Figure 3E), no significant group differences were apparent (*F*(2,33) = 1.42, *p* = 0.26).

**Figure 3.**
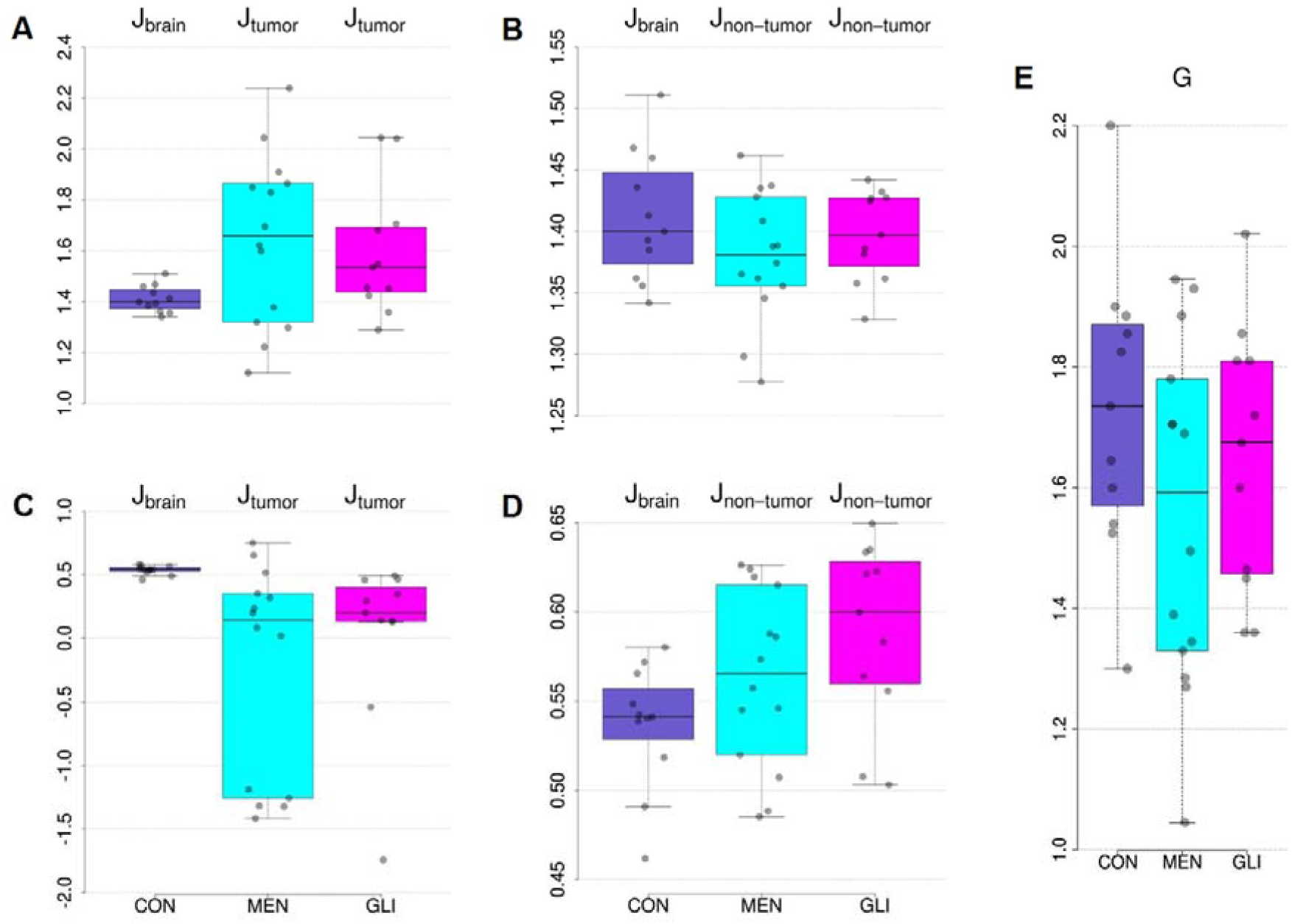
Distributions of optimal modeling parameter values per group of (A) median local inhibitory connection strengths across entire brain in healthy controls, and across tumor regions in meningioma and glioma patients; (B) median local inhibitory connection strengths across entire brain in healthy controls, and across nontumor regions in meningioma and glioma patients; (C) median local inhibitory connection strengths, corrected for region size, across entire brain in healthy controls, and across tumor regions in meningioma and glioma patients; (D) median local inhibitory connection strengths, corrected for region size, across entire brain in healthy controls, and across non-tumor regions in meningioma and glioma patients; and (E) global scaling parameter. CON = healthy control participant; MEN = meningioma patient; GLI = glioma patient.

### Comparison of structural network topology and cognitive performance between brain tumor patients and controls

Before relating the biophysical model parameters to cognitive performance and structural network topology, we examined whether group differences were present in these variables. As can be seen in Figure 4A, results of one-way ANOVA tests showed no significant group differences between glioma patients, meningioma patients and healthy controls in any of the cognitive domains assessed (*F*(2,32) = 0.13, *p* = 0.88 for reaction time; *F*(2,31) = 0.35, *p* = 0.71 for sustained attention; *F*(2,32) = 0.17, *p* = 0.84 for planning accuracy; *F*(2,32) = 0.38, *p* = 0.69 for spatial span length). With regard to structural network topology, a significant group difference was found in the participation coefficient (*F*(2,33) = 3.94, *p* = 0.029) (Figure 4B, right). Post hoc (Tukey) testing revealed that participation coefficient was higher in glioma patients compared to healthy controls (*p* = 0.026). Group differences in global efficiency and modularity were not significant (*F*(2,33) = 0.57, *p* = 0.57 for global efficiency; *F*(2,33) = 1.26, *p* = 0.30 for modularity) (Figure 4B, left & middle).

**Figure 4.**
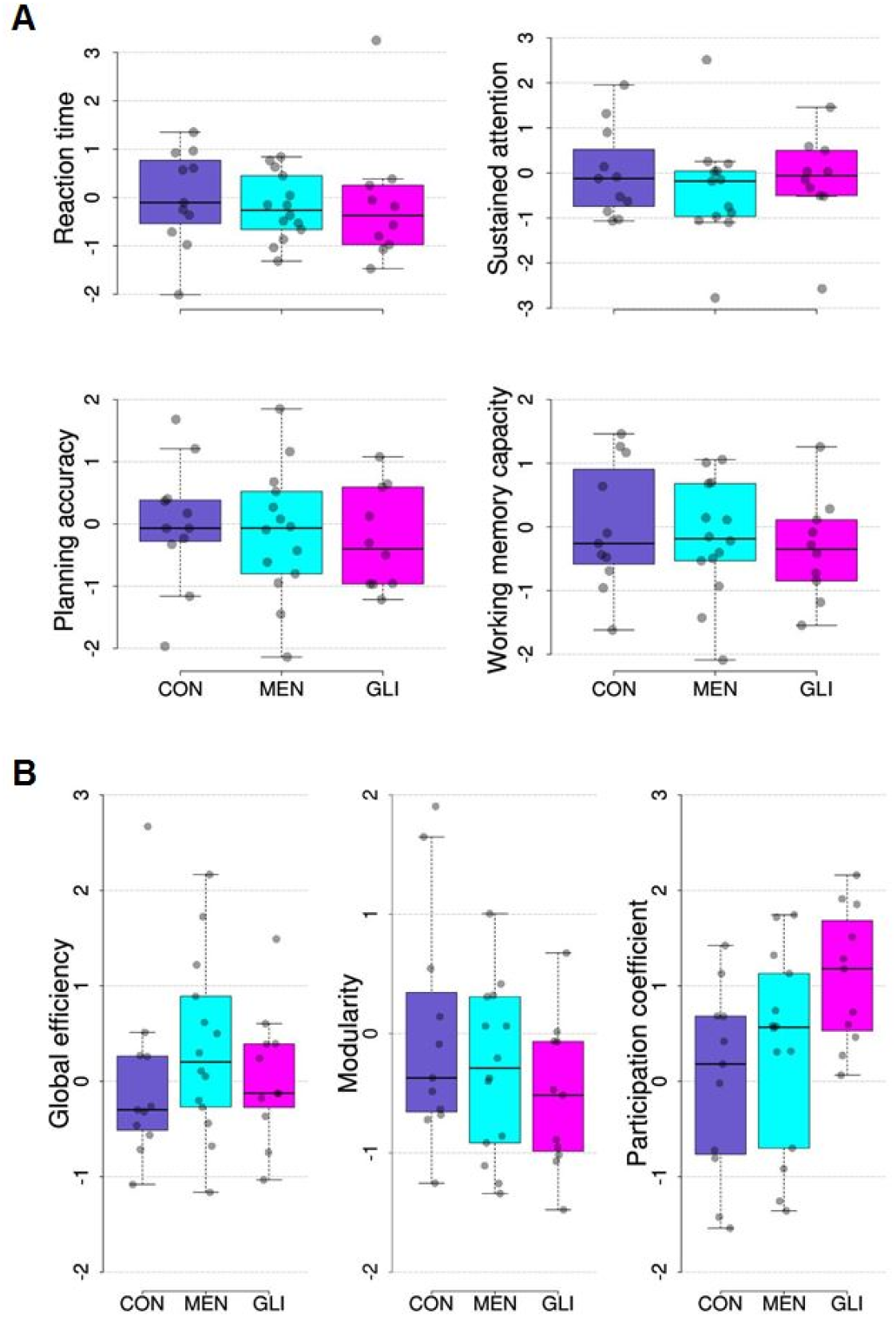
Distribution of (A) cognitive performance measures and (B) graph theory metrics per group. All metrics are corrected for important confounding variables (described in methods section) and transformed to z-scores using the mean and standard deviation of the respective metric in the group of control subjects. CON = healthy control participants, MEN = meningioma patients, GLI = glioma patients.

### Modeling parameters are associated with structural network topology and cognitive performance

Next, we aimed to elucidate the relationship between the individually optimized modeling parameters on the one hand, and structural network topology and cognitive performance on the other hand. Linear regression analysis showed that global efficiency of the structural network was significantly associated with the global scaling factor (*t* = −4.42, *p* < 0.001, *η^2^* = 0.34) and the median feedback inhibition control parameter across non-tumor regions in brain tumor patients (corrected for region size) (*t* = −2.67, *p* = 0.014, *η^2^* = 0.25). Specifically, higher values of global efficiency were associated with lower values of both modeling parameters, as illustrated in Figure 5 (Pearson’s *r* = −0.67 and −0.45 for the global coupling parameter and the feedback inhibition control parameter, respectively). No significant associations were found between the modeling parameters and modularity or participation coefficient.

**Figure 5.**
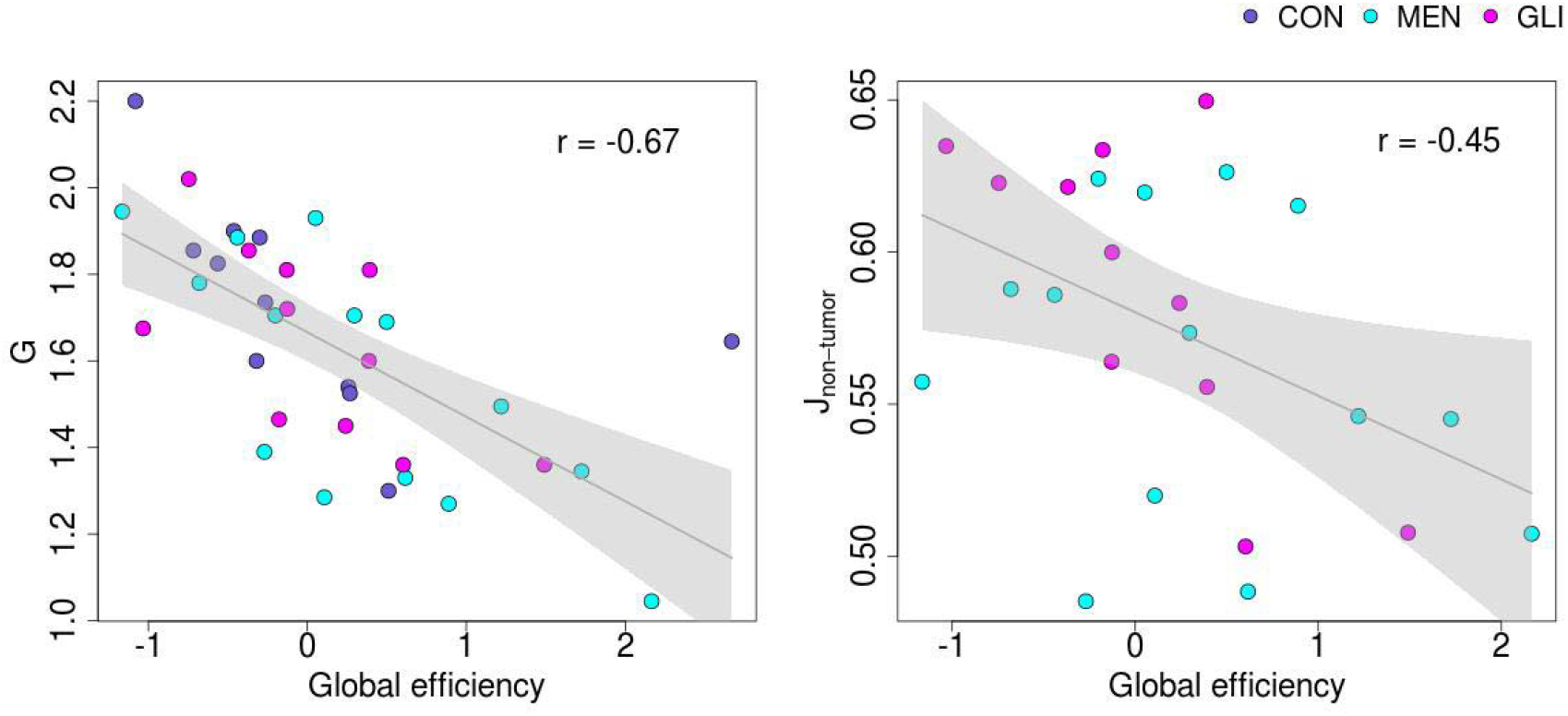
Linear relationship between global efficiency on the one hand and global coupling parameter (left) and median feedback inhibition control parameter across non-tumor regions in brain tumor patients (corrected for region size) (right) on the other hand. Gray line represents regression line with 95% confidence interval. Group membership is color-coded: CON = healthy control participants; MEN = meningioma patients; GLI = glioma patients.

Furthermore, a significant negative association was found between the median feedback inhibition control parameter across tumor regions (corrected for region size) and patients’ reaction time (*t* = −4.23, *p* = 0.001, *η^2^* = 0.44) and sustained attention (*t* = −2.62, *p* = 0.018, *η^2^* = 0.17) (Figure 6, left). In contrast, these associations were absent in non-tumor regions (*t* = 1.27, *p* = 0.222, *η^2^* = 0.07 for reaction time; *t* = 0.61, *p* = 0.549, *η^2^* = 0.02 for sustained attention) (Figure 6, right). However, as can be seen in Figure 6, the correlation between local inhibitory connection strengths and sustained attention is mainly driven by a few outlying observations, hence caution is advised in interpreting this finding. No significant associations were found between individual modeling parameters and working memory capacity or spatial span length.

**Figure 6.**
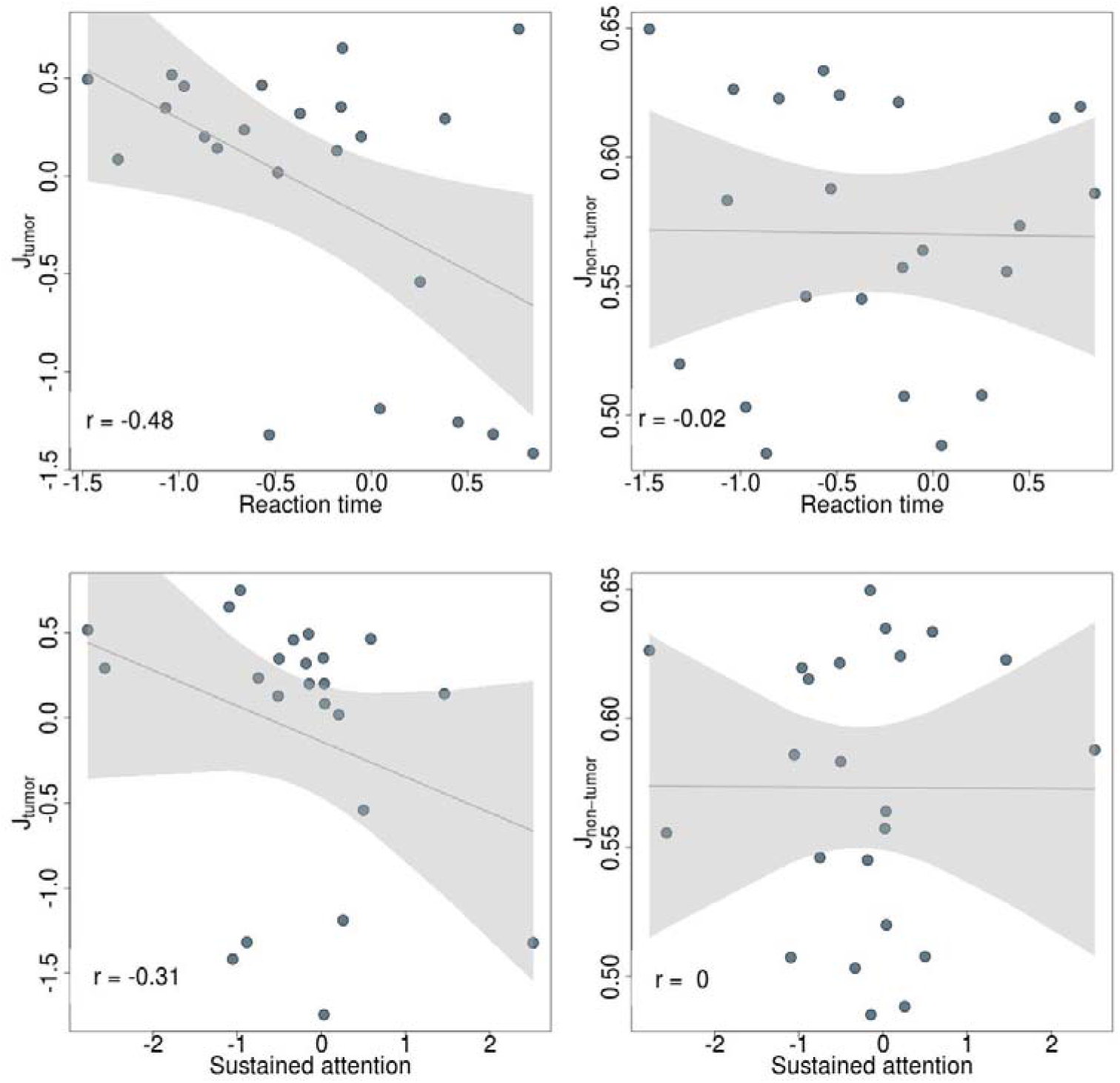
Linear relationship between reaction time (top) and sustained attention (bottom) and feedback inhibition control parameter (corrected for region size) across tumor (left) and non-tumor (right) regions in meningioma and glioma patients. Gray line represents regression line with 95% confidence interval.

## Discussion

Applying The Virtual Brain to simulate large-scale brain dynamics in brain tumor patients and healthy controls, we have demonstrated (1) that utilizing personalized virtual brain models significantly improve prediction accuracy of simulated functional connectivity; (2) that local inhibitory connection weights can differentiate between regions directly affected by a brain tumor, regions more distant from a brain tumor, and regions in a healthy brain; and (3) that individually optimized model parameters correlate with structural network topology and cognitive performance.

### The importance of personalized virtual brain models

Individualized medicine is increasingly put forward as an important means to advance medical care. In this regard, The Virtual Brain holds great promise, given its direct focus on the simulation of subject-specific brain dynamics. Reliable prediction of patient-specific large-scale brain dynamics would open up the possibility to virtually lesion structural connectomes, making computational models unique predictive tools to investigate the impact of diverse structural connectivity alterations on brain functioning. That is, computational modeling would enable us to investigate what types or extent of damage the brain can withstand, and conversely, which kind of distortions can be expected after brain lesions, including those purposively induced by surgery.

Therefore, the first aim of this study was to verify the added value of personalized virtual brain models. Results showed that individual optimization of the modeling parameters significantly increased the prediction accuracy of individual functional connectivity. However, whether model parameters were tuned on the individual SC matrices or on the average SC matrix across all control subjects did not yield a significant difference, corroborating findings in previous research (e.g., Jirsa et al., 2017). Future computational modeling studies could utilize finer and more elaborated parcellation schemes, for example based on multimodal data (Glasser et al., 2016), to investigate whether individual traits can be captured through tractography that have relevance for the prediction of individual functional connectivity.

### Predictors of individual biophysical model parameters

In the next step, we investigated which factors are associated with the individual model parameters, examining the relative contribution of the presence and type of brain tumor, structural network topology, and cognitive performance.

With regard to the effect of the presence and type of brain tumor, results revealed no significant differences in the feedback inhibition control parameter between tumor regions and healthy regions/brains. However, this local parameter is highly dependent on the size of the cortical area that is modeled, and additional analyses showed that regions affected by a tumor were on average larger than non-tumor regions. After taking into account this confounding effect of region size, results showed median local inhibitory connection strengths that are much lower and more variable in tumor regions compared to healthy regions/brains. Moreover, local inhibitory connection strengths (controlled for region size) in non-tumor regions were increased compared to healthy controls. Hence, these results suggest that brain tumors have a distinct focal and distant effect on the local inhibitory connection weights, beyond what could be expected from their region size.

A previous computational modeling study also reported alterations in inhibitory over excitatory coupling in chronic stroke patients (Falcon et al., 2016). It is however difficult to compare these results to those of our study, as Falcon and colleagues did not correct for region size or differentiate between affected and unaffected cortical areas. At the microlevel, seminal studies by Sontheimer and colleagues (Buckingham et al., 2011; Sontheimer, 2008) have demonstrated that gliomas cause excessive peritumoral glutamate release; a major excitatory neurotransmitter in the brain. This excessive release leads to local hyperexcitability in the vicinity of the tumor, disturbing the delicate excitation/inhibition balance, which in turn has been found to cause epileptic activity in glioma patients. Future research, using neuroimaging data with a much finer spatial resolution, would however be needed to investigate whether these cellular alterations in excitation/inhibition balance can be detected by the feedback inhibition control model parameter in regions close to glioma tumors.

In contrast to previous computational modeling studies that found increased global coupling values in chronic stroke patients (Falcon et al., 2015, 2016), our results revealed no significant differences in global coupling between glioma patients, meningioma patients and healthy controls.

We then turned to structural network properties and measures of cognitive performance as possible predictors of the individual biophysical model parameters. Initial descriptive analyses of these metrics revealed no significant differences in cognitive performance between brain tumor patients and healthy controls. Although most patients with slow-growing brain tumors exhibit normal clinical exams (Walker & Kaye, 2003), previous studies have found slight cognitive deficits using extensive assessments (for a review see Taphoorn & Klein, 2004). Regarding structural network topology, results showed no significant group differences in global efficiency and modularity, corroborating a previous study in which structural network integration and segregation properties were found to be mostly preserved in brain tumor patients, compared to healthy controls (Yu et al., 2016). We did however find a significant increase in participation coefficient in glioma patients compared to healthy controls. The observed higher participation coefficients imply that links in the structural network of glioma patients are on average more uniformly distributed among distinct modules, which may facilitate communication or integration of multiple types of information (Power, Schlaggar, Lessov-Schlaggar, & Petersen, 2013). Although this finding warrants further investigation, we hypothesize that focal brain damage might have stimulated rewiring of affected nodes to neighboring communities in order to preserve functionality. This could then also have prevented alterations in network segregation and integration.

Despite a lack of large group differences in cognitive performance and structural network topology, several interesting associations across groups were found between these variables and the individually optimized biophysical model parameters. First, results showed a strong negative relation between global efficiency of the structural network and global coupling, corroborating the finding of Falcon et al. (2015) in stroke patients. This implies that higher global coupling values are required in subjects whose structural connectome is less efficiently organized, in order to achieve the same amount of functional connectivity between cortical areas. In addition, decreased global efficiency was found to be related to increased inhibition of excitatory neuronal populations in regions distant from the tumor, but *not* in those directly affected by the tumor. Put differently, when brain tumor patients’ structural network is less efficiently organized, it requires not only higher global coupling but also more local inhibition in regions that are not directly affected by the tumor, to achieve the same amount of synchronization. This finding demonstrates the widespread impact of brain tumors, despite their relatively focal damage, and hence could be termed “modeled diaschisis” as an extension to the concepts of “connectional diaschisis” and “connectomal diaschisis” (Carrera & Tononi, 2014).

Furthermore, negative associations were found between local inhibitory connection weights in regions directly affected by a brain tumor and reaction time and sustained attention. This implies that brain tumor patients who performed worse on sustained attention and reaction time tasks on average had higher local inhibitory connection weights in regions directly affected by the tumor. This finding is a bit counterintuitive, since the feedback inhibition control parameter was significantly lower in brain tumor patients compared to healthy controls, which would imply a higher cognitive performance in brain tumor patients relative to healthy controls. However, as noted in the results section, the correlation between local inhibitory connection strength and especially sustained attention is mainly driven by a few outlying observations, hence caution is advised in interpreting this finding. Future research using a larger sample size would be required to clarify this association.

### Limitations and future directions

In interpreting the results of this study, some important limitations should be taken into consideration. First, the sample size is rather small, limiting the statistical power of the analyses. In addition, substantial inter-subject variability is present in both patient groups, caused by (among other factors) heterogeneity in lesion etiology and size. It is likely that these two factors interfere with separating clinical groups based on computational model parameters, cognitive performance and structural network topology. As more efforts are undertaken to make clinical datasets publicly available, future studies would benefit from using larger sample sizes.

Second, simulated and empirical functional connectivity were only moderately related after optimization of the model parameters. Moreover, using individual structural connectomes did not yield better predictions of individual functional connectivity patterns compared to using a control average structural connectivity matrix. This is, however, a limitation of all current computational modeling studies and a matter of much debate. A recent study by Zimmermann and colleagues (Zimmermann, Griffiths, Schirner, Ritter, & McIntosh, 2018) focused on this exact issue and reported that subject-specificity of SC-FC is limited due to the relatively small variability between subjects in SC compared to the larger variability in FC. This limited variability in individual SC matrices could be due to the quality of current individual SC matrices. Although great advances have been made in diffusion weighted imaging acquisitions and tractography algorithms, it is known that DWI tractography underestimates the number of short-distance streamlines in favor of long fiber tracks (Jeurissen et al., 2017). Although in the current study we have used a relatively new multi-shell DWI sequence with b-values of up to 2800 s/mm^2^ and a state-of-the-art processing pipeline, future studies could investigate whether prediction accuracies of individual SC matrices improve when using data of even better quality (using for example 7T MRI scanners, with multiband sequences and/or longer acquisition times). Another contributing factor to the low variability in individual SC matrices might be the relatively coarse parcellation schemes currently applied in computational modeling studies. With increasing computational power, future studies can test finer and more elaborate parcellations, for example based on multimodal data such as in Glasser et al., (2016) to investigate whether individual traits can be captured through tractography that have relevance for the prediction of individual FC.

Related to this issue, the robustness of the obtained results to different parameter space exploration criteria has to be investigated. In this study, the link-wise Pearson correlation between empirical and simulated FC was maximized. Though this method is routinely employed in large-scale modeling studies, other methods are worth exploring. For example, similarity could be sought at the modular level, maximizing the cross-modularity between simulated and empirical functional connectivity (Diez et al., 2015; Stramaglia et al., 2017). These approaches have the additional advantage of reducing the dependency on the parcellation choice.

As pointed out in the introduction, the next step entails prediction of post-surgical outcome. To this end, longitudinal research is required to investigate whether individually optimized model parameters can reliably predict post-surgical functional connectivity. This would be a major step towards pre-surgical virtual exploration of different neurosurgical approaches and to identify an optimal surgical strategy.

**Extended Data 1.** Code used for simulation of large-scale brain dynamics and scripts for subsequent postprocessing analyses.

